# Quantitative photoacoustic imaging study of tumours *in vivo*: baseline variations in quantitative measurements

**DOI:** 10.1101/307595

**Authors:** Márcia Martinho Costa, Anant Shah, Ian Rivens, Carol Box, Tuathan O’Shea, Efthymia Papaevangelou, Jeffrey Bamber, Gail ter Haar

## Abstract

Photoacoustic imaging (PAI) provides information on haemoglobin levels and blood oxygenation (sO_2_). To facilitate assessment of the variability in sO_2_ and haemoglobin in tumours, for example in response to therapies, the baseline variability of these parameters was evaluated in subcutaneous head and neck tumours in mice, using a PAI system (MSOT-inVision-256TF). Tumours of anaesthetized animals (midazolam-fentanyl-medetomidine) were imaged for 75 minutes; in varying positions; and repeatedly over 6 days. An increasing linear trend for average tumoural haemoglobin and blood sO_2_ was observed, when imaging over 75 minutes. There were no significant differences in these temporal trends, when re-positioning tumours. A negative correlation was found between the percent decrease in blood sO_2_ over 6 days and tumour growth rate. This paper shows the potential of PAI to provide baseline data for assessing the significance of intra- and inter-tumoural variations that may eventually have value for predicting and/or monitoring cancer treatment response.

## 1. Introduction

The immature, tortuous and hyper-permeable vascular network often found in tumours may not provide sufficient nutrients and oxygen for rapid and uncontrolled cell proliferation, which can result in regions of tumour cell hypoxia, i.e. low oxygen concentration in the vicinity of tumour cells [1]. Temporal fluctuations in the hypoxic state also occur [2], on a time-scale of minutes or hours when they result from an efflux of red blood cells, or days when they are due to vascular network remodelling and angiogenesis [3].

Hypoxia is associated with increased resistance to chemo- and radio-therapy (as extensively reviewed by Rockwell et al. [4]), two commonly used clinical cancer treatments. Assessing the spatial and temporal variation in hypoxia levels could be predictive of a cancer therapy response and may therefore play a critical role in treatment planning and adaptation [5, 6]. Conventionally, oxygen needle electrodes have been used to assess hypoxia in patients, by directly measuring tissue partial oxygen pressure (pO_2_) [7]. However, needle electrodes are invasive and provide only a local value, which is not representative of the whole tumour. Techniques which use conventional non-invasive imaging have been developed to attempt to infer, not only the oxygenation levels, but also the spatial distribution of hypoxia. Several Positron Emission Tomography (PET) radiotracers are available [8–10] which are molecules that are reduced in the absence of oxygen and covalently bind to intracellular molecules, providing a non-invasive alternative, albeit with a relatively poor spatial resolution and high cost. Unfortunately, the use of radioactive isotopes is also ill-suited to longitudinal studies due to radiation dose accumulation. Blood oxygenation level dependent (BOLD) magnetic resonance imaging (MRI) [11] enables the measurement of relative blood oxygen saturation (sO_2_) increase during carbogen challenge compared to air-breathing, within tumours. It has been shown that these measurements are in good spatial agreement with tissue pO_2_ measurements [12–14] and histological markers of hypoxia [15, 16]. BOLD-MRI, however, has poor specificity for hypoxia, and does not provide absolute sO_2_.

Recently, the use of combined ultrasound (US) and optical imaging techniques, such as photoacoustic imaging (PAI), also known as optoacoustic imaging, has been proposed as a method of non-invasively mapping blood oxygen saturation [17, 18]. PAI combines the merits of optical and ultrasound imaging, displaying optical absorption contrast with sub-millimetre spatial resolution at depths of several centimetres in tissue. Light is absorbed differentially depending on the relative concentrations of light-absorbing molecules present [19]. As a result, PAI can identify the unique spectral signatures of deoxy-(Hb) and oxy-haemoglobin (HbO_2_) [17]. The quantification of the local levels of each chromophore further allows mapping of blood sO_2_ distribution in the tissue, defined as the ratio of oxygenated haemoglobin (HbO_2_) to the total amount of haemoglobin (HbT, the sum of HbO_2_ and deoxy-haemoglobin, Hb) [19, 20]. Low sO_2_ regions tend to be hypoxic [21].

PAI has been used for hypoxia mapping pre-clinically, in subcutaneous [15, 18, 20, 22–25] and brain tumours [26], showing good agreement with pimonidazole staining [18, 22], an exogenous hypoxia marker [15], and imaging modalities, such as bioluminescence [20], high resolution and dynamic contrast enhanced ultrasound [20, 23] and BOLD-MRI [24, 25].

This paper aims to characterise the variability of photoacoustic imaging measurements of Hb, HbO_2_, HbT and sO_2_ in subcutaneous head and neck tumours, commonly hypoxic, using a commercial tomographic photoacoustic system. The long term application for this study would be to use the baseline measurements obtained here, in tumour-bearing control animals, in future longitudinal imaging studies where two cancer therapies will be used, radiotherapy and high intensity focused ultrasound (HIFU). This baseline information is valuable to assess the significance of any difference in pre-treatment blood sO_2_ or haemoglobin levels from the mean across the tumours and any effect of the treatments on these parameters, as it has been reported in the literature that both these therapies can alter the vasculature and/or the oxygenation status of tumours after being delivered [27–29]. Therefore, this study aims to obtain baseline measurements of the variation in the photoacoustic imaging measurements over the short term (<75 minutes), to investigate the potential effects of both anaesthetic duration and altering the position of the animal, as well as over the longer term (6 days), to evaluate the variations in growing tumours.

## 2. Methods

### 2.1. Animal experiments

All research was conducted under the Guidelines of Animal Welfare provided by the UK Home Office with approval from the local animal welfare and ethics board and under Helsinki Declaration of 1975.

The human head and neck squamous cell tumour model, CAL^R^, used for this study is a non-commercially available cancer cell line developed at the Institute of Cancer Research [30]. Half a million cells were injected subcutaneously into the right flank of 6 week old female FOXnu^n1^ mice (˜25 g). Animals were monitored for the development of a mass. Once palpable, tumour dimensions were measured every 2 or 3 days, in 3 orthogonal directions (length (l), width (w) and height (h)), using Vernier calipers. Tumour volume (V) was calculated assuming an ellipsoidal shape, commonly used for subcutaneous tumours, according to Eq. 1.

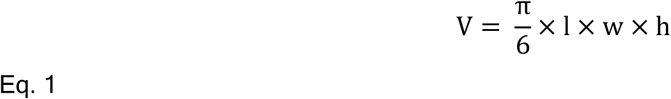

The volume error (V_error_), for each individual tumour, was estimated from the derivative of an ellipsoidal volume, assuming a measurement error (E_m_) of 0.5 mm in each direction, using Eq. 2.

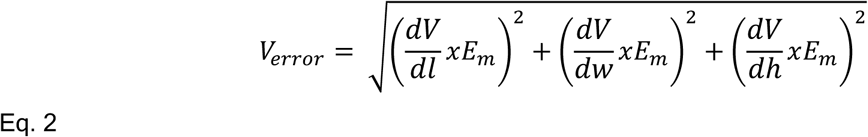

Tumour dimensions are therefore presented as mean volume plus standard deviation of the estimate based only on the accuracy of the three orthogonal dimension measurements, ignoring any deviation in shape from an ellipsoid.

### 2.2. Tumour growth rates

Tumour growth curves were fitted to the Exponential-Linear Model to calculate tumours’ growth rates [31]. The model assumes that cells initially proliferate with constant cell cycle duration, T_c_, resulting in exponential growth, followed by a linear growth phase, when there is a decrease in actively proliferating cells and these are constrained to the margins of the tumour [32]. The model is described as follows:

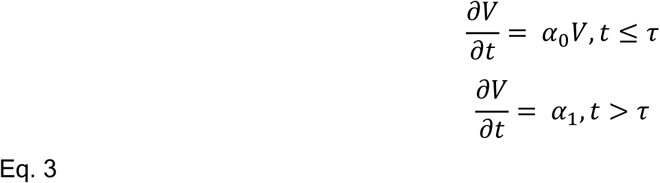

where V is tumour volume (mm^3^), t is time (days), α_0_ is the growth rate during the exponential phase, i.e. the fraction of proliferating cells at time ln2/T_c_, α_1_ is the growth rate in the linear phase and τ represents the time point at which the growth changes from the exponential to the linear phase. Assuming the solution is continuously differentiable then:

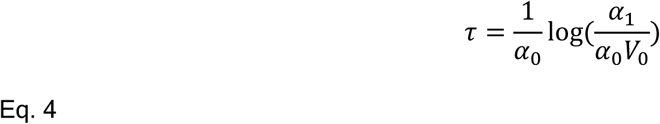

where V_0_ is the initial volume, in this case 0.1 mm ^3^, chosen as an arbitrary small initial volume of tumour cells.

The tumour growth fitting was carried out in MATLAB (2013a, 8.1.0.604), using a nonlinear least squared fit to obtain the α_0_ and α_1_ parameters. Fits were performed on volume data weighted using the measurement uncertainties of each data point, which corresponded to the V_error_ calculated from Eq. 2, within 95% confidence interval bounds.

### 2.3. Photoacoustic imaging data acquisition and reconstruction

Mice were imaged using a commercially available, real time, multispectral optoacoustic tomographic (MSOT) device, inVision 256-TF small animal scanner (iThera Medical GmbH, Munich, Germany) [33]. The system has a tunable optical parametric oscillator pumped by a Q-switched Nd:YAG near-infrared laser. A 256 element, toroidally focused, ultrasound imaging transducer (4 cm radius) with a centre frequency of 5 MHz (60% bandwidth) was used to acquire the acoustic signal, resulting from the optical tissue excitation. These elements cover an angle of 270 degrees around the tumour to create a cross-sectional image.

Tumours were first imaged (defined as Day1) when they reached a volume of approximately 200 mm^3^. Anaesthesia was induced using an intraperitoneal injection of midazolam:fentanyl:medetomidine (5.0:0.05:0.5 mg.Kg^−1^). Mice were placed horizontally in a holder encased in a thin (17 µm) polyvinyl chloride (PVC) film (Fig.1), with their tumour facing vertically downwards, i.e. directly in the direction of the laser beam, and acoustically coupled to the membrane using ultrasound imaging gel. In order to provide acoustic coupling to the MSOT, the animal holder was completely submerged in a tank filled with 34°C distilled water. Medical air (21% oxygen) and/or 100%-oxygen were supplied continuously to the mouse via tubing during immersion in order to avoid asphyxiation. Imaging started approximately 5 minutes after placing the holder in the water tank, in order to allow the animal’s body temperature to equilibrate with the water’s temperature.

**Fig. 1.**
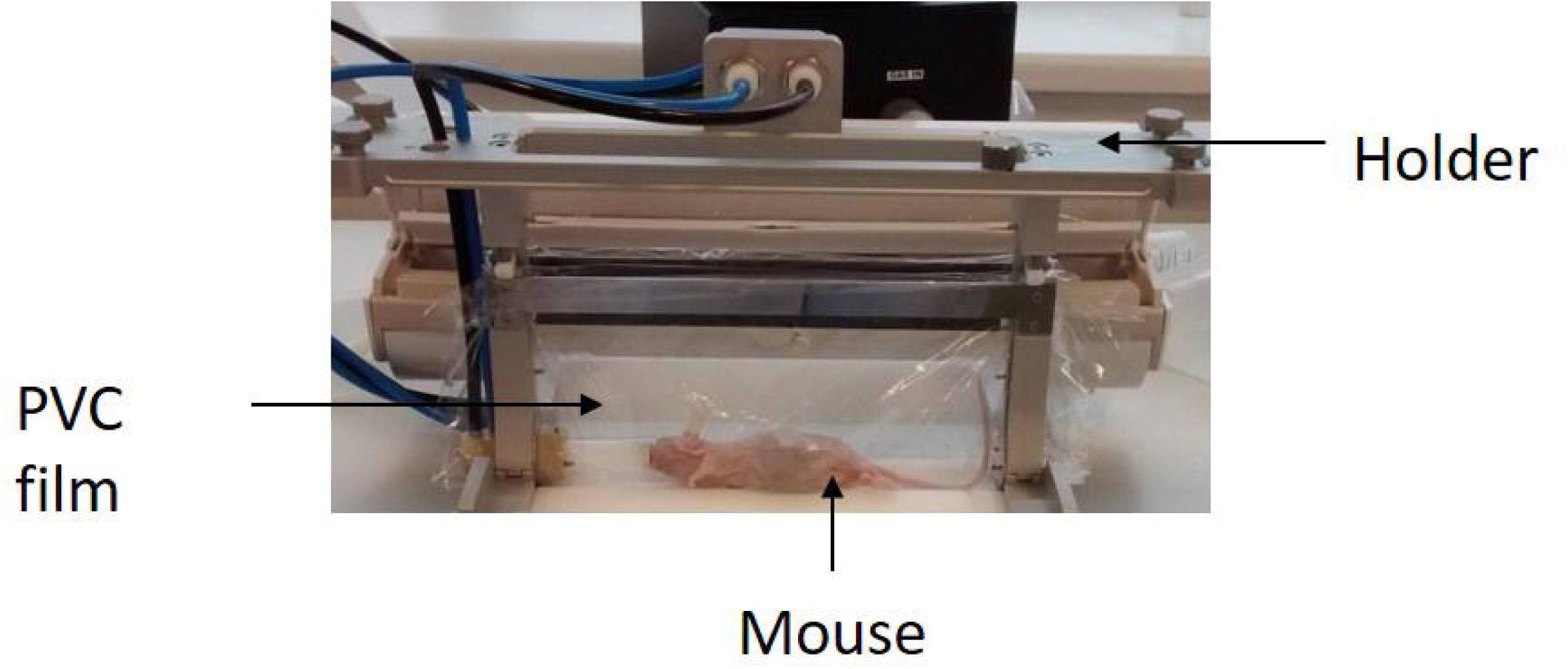
Photograph showing the experimental setup of the mouse, in the MSOT holder prior to mounting in the imaging system.

For acquiring the photoacoustic data, a volume of interest (VOI), containing the whole tumour was selected to avoid a large dataset. Transverse (medial-lateral direction) imaging slices were acquired at 1 mm intervals in the cranial-caudal direction. The optical imaging wavelengths used were: 700, 715, 730, 750, 760, 800, 850, and 900 nm. This wavelength range maximises the differences in Hb and HbO_2_ signals. Ten signal averages per wavelength were obtained for each imaging slice.

Photoacoustic images were reconstructed using a model-based inversion algorithm [34] which was implemented in the viewMSOT software (v3.8) provided by iThera Medical. Spectral unmixing was used to calculate the contribution of each chromophore, oxy- and deoxy-haemoglobin, in blood, in order to calculate the oxygen saturation. This is done on a pixel-by-pixel basis, through a linear regression algorithm [35]. The software subsequently allows calculation of the mean oxy- (HbO_2_) and deoxy-haemoglobin (Hb) signals in a user defined region of interest (ROI) which could be drawn onto the greyscale anatomical image (Fig. 2A). These mean Hb and HbO_2_ values are added together to calculate the mean total haemoglobin per ROI. For any ROI the mean oxygen saturation was calculated using Eq. 5.

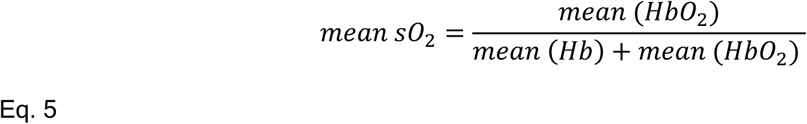

**Fig. 2.**
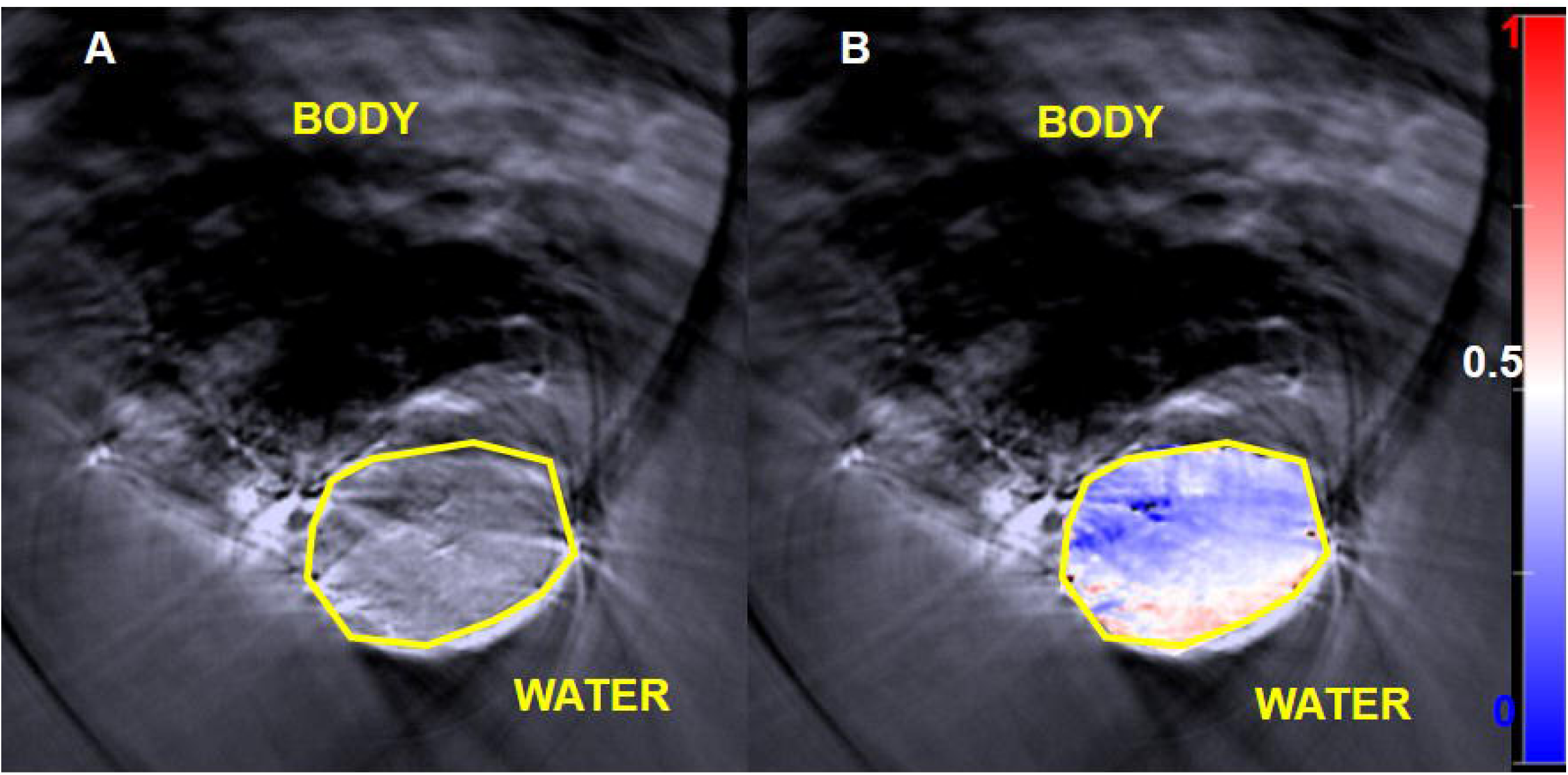
A: Greyscale anatomical photoacoustic image obtained from optical excitation at a single wavelength (800 nm). The yellow polygon is the ROI used to delineate the tumour. B: Shows an ‘oxymap’ overlaid on the tumour. The ‘oxymap’ colour scale on the right represents the blood sO_2_ calculated by the MSOT software on a pixel-by-pixel basis. Blue represents sO_2_ <0.5 A.U (white), while red is sO_2_ >0.5 A.U. Black pixels within the ROI show where haemoglobin signals are below the noise threshold.

All the parameters are unitless quantities, described in arbitrary units (A.U.), because the software does not take use light fluence and the system response in calculations, which would be necessary to quantify the concentration (in mmol/L or mg/L) of each chromophore. For each 3D imaging scan, 3 slices were always chosen for the analysis of Hb, HbO_2_, HbT and sO_2_.

The viewMSOT software also allows mapping of the spatial distribution of sO_2_ calculated on a pixel-by-pixel basis, an ‘oxymap’ (Fig. 2B). Black pixels in such images indicate where the total amount of haemoglobin is so low, or sources of noise so high, that adequate confidence in spectral recognition cannot be achieved, or because the reconstruction method produces pixels with negative values. The intensity of such pixels is set to zero. The software automatically provides the percentage of black pixels per ROI.

### 2.4. Imaging studies

#### Three separate imaging studies were performed

The first was a short term MSOT study designed to investigate any potential effects of anaesthesia on total haemoglobin and blood sO_2._ A 3D whole tumour dataset was acquired every 5 minutes for 75 minutes after allowing 5 minutes for the tumour to reach thermal equilibrium with the imaging system. Only air-breathing imaging was acquired, in order to maximise the rate (and number) of datasets that could be acquired. Measurements were acquired in 4 animals when their tumours reached a target volume of 200 mm^3^.

The PAI optical fluence is depth-dependent, hence the position of tissue of interest with respect to the imaging transducer can affect the PAI measurements, as this can alter the amount of near-infrared light attenuated by overlying tissue as well as the field intensity incident on that position. In order to study how re-positioning the animal affects measurements of blood haemoglobin and blood sO_2_, a second short term study was performed, 24 hours after the first study. The same 4 animals were anaesthetised once and imaged in three consecutive sessions after being dismounted from the imaging system between each session. In addition, the tumour was placed at a slightly different position relative to the transducer each time. Whole tumour 3D datasets were acquired first under air breathing conditions which took approximately 2 minutes, and then under 100% oxygen breathing, after allowing 2 minutes for the oxygen to reach the tumour, for the same VOI, under hyperoxic conditions [25]. Oxygen breathing can be used as a contrast agent for PAI, enhancing tumour and spatial differences when vascularity is imaged using sO_2_ [36]. Approximately 4 minutes were needed to remove the animal from the holder, reposition it and replace it (see Fig. 3).

**Fig. 3.**
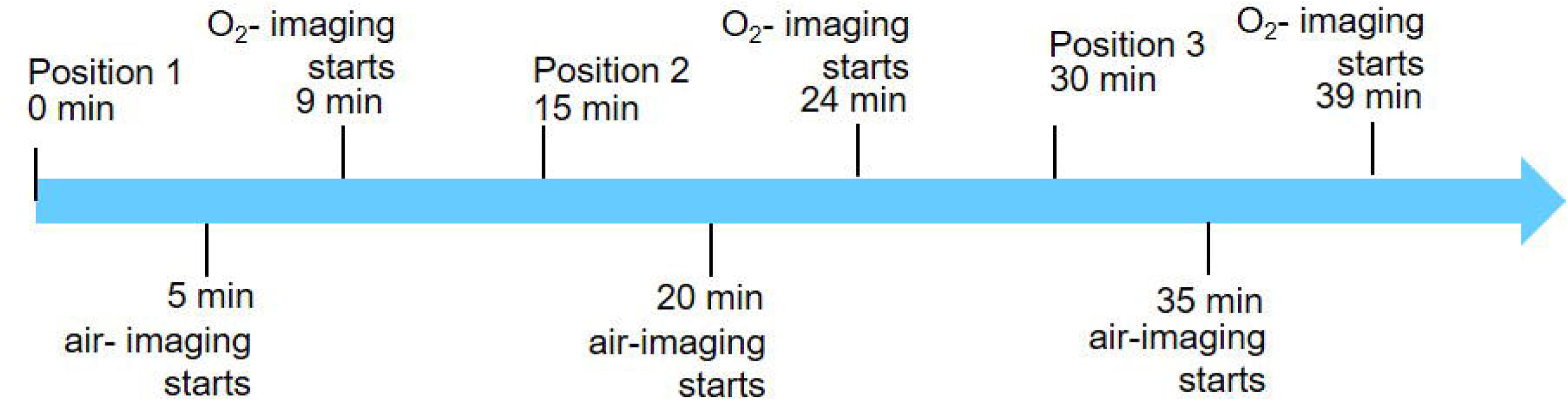
Diagram for the re-positioning of the animal. Each air-breathing imaging session started at approximately 5, 20 and 35 minutes after placing the animal in the water tank of the MSOT system. Air-breathing imaging takes approximately 2 minutes. The air is then changed for 100 %-O_2_, for 2 minutes and the same tumour region is re-imaged for approximately 2 minutes again, during O_2_-breathing.

For both short-term studies, a linear regression model was applied to calculate an estimate of the average change in signal over time, as well as the goodness of fit between the imaging data and the model. This was obtained using the curve fitting toolbox (version 3.5.3) available in MATLAB R2016a software. To investigate the suitability of the linear model for these studies, three parameters were calculated: goodness of fit (R^2^) that indicates how much of the variability of the parameter (Hb, HbO_2_, HbT and sO_2_) is time-dependent; p-value of the model represents if the slope is significantly different from 0 (if p-value<0.05); and the Sum of Squared Errors (SSE) of the prediction model that indicates the suitability of the linear model to explain the variability behaviour of the data (the closer SSE is to 0, the better the linear fitting). The temporal variations of both 75 minute and re-positioning study were also compared, by calculating if there were statistical differences in signal change for both studies, using an analysis of covariance (software: GraphPad Prism 7).

The third, a long-term study, was designed to sequentially image tumours during a period of growth. When tumours (n=10) reached a volume of approximately 200 mm^3^, animals were imaged on 3 consecutive days (‘Day 1, 2 or 3’). This volume was chosen as being suitable for future studies of cancer treatment e.g. using radiotherapy. Tumours were also imaged on day 6 in order to evaluate changes in haemoglobin and blood sO_2_ during tumour growth. For this study, animals were imaged under air- and oxygen-breathing conditions.

For short- and long-term studies, three slices in each tumour, separated by 1 mm, were chosen for analysis, a central slice and the adjacent slice on either side. The mean haemoglobin and blood sO_2_ values were obtained by averaging over the three whole-tumour ROIs to provide the tumour ROI-averaged Hb, HbO_2_, HbT and sO_2_. For each animal, the average percentage of black pixels in the three central tumour slices was also calculated in order to evaluate the signal lost in each ROI.

### 2.5. Immunohistochemistry

Five tumours were collected after imaging for immunohistochemistry, including pimonidazole staining that binds to hypoxic ducts, and H&E to analyse tissue structure. Pimonidazole was administered to the animal intraperitoneally (60 mg/kg) 45 minutes before sacrificing it. Tumour samples were excised and immediately snap frozen on dry ice and stored at −80°C until sectioning.

For H&E staining, frozen tumour sections (10 µm thick) were counterstained in ‘Harris‘ haematoxylin (dyes cellular nuclei) and eosin (dyes cytoplasm and organelles). Samples were then dehydrated with a sequence of ethanols and xylene, and mounted using DPX mounting medium (Sigma-Aldrich, Dorset, UK). Acetone-fixed cryosections (consecutive to the ones used for H&E) were used to visualize pimonidazole adduct formation with an anti-pimonidazole FITC-conjugated mouse monoclonal antibody (1:200) (Hypoxyprobe™ Plus kit, Hypoxyprobe, Inc., Burlington, USA).

Stained sections were imaged using a BX51 microscope (Olympus Optical, London, UK), with a motorised stage (Prior Scientific Instruments, Cambridge, UK), driven by microimaging software (cellSens, Olympus Optical, London, UK). This allows automatic acquisition of the full sample. Microscopic images were acquired using a digital camera (ColorView 12; Soft Imaging System GmbH, Munster, Germany) and exposure times of 100 ms (H & E) or 500 ms (pimonidazole) using a x4 objective.

### 2.6. Statistical analysis

The slice-, intra-tumour and inter-tumour coefficients of variation (CoVs), i.e. the ratio of standard deviation to mean, were calculated for Hb, HbO_2_, HbT and sO_2_ in all PAI studies. The first indicates how much variation to expect between the 3 ROIs chosen for the analysis of the PAI parameters; the second the variation of the measurements for each animal and the third the variation between animals.

A paired student t-test was used to assess the level/degree of significance of differences in the mean Hb, HbO_2_, HbT and sO_2_ of CAL^R^ tumours during the long term variability studies, on imaging days 1, 2, 3 and 6, using R software.

## 3. Results

### 3.1. Comparison between photoacoustic imaging and immunohistochemistry

Pimonidazole staining indicates the presence of widespread perfusion-limited or diffusion-limited hypoxia (Fig.4B and E) with unstained regions largely corresponding to necrotic/acellular regions, observed in the H&E staining (Fig. 4A & D) and avascular (black or very dark blue) regions on the PAI ‘oxymaps’. The blue pixels in tumour ‘oxymaps’ (Fig. 4C and F) corresponding to low sO_2_ (<0.5), spatially correlated to tumour hypoxia, as determined using pimonidazole staining. Red pixels in the tumour ‘oxymaps’, e.g. the periphery of the tumour in Fig. 4C, generally corresponded with weak/negative pimonidazole staining (Fig. 4B). Albeit, a small proportion of the red oxymap pixels (particularly in Fig. 4C), correspond with pimonidazole staining, indicating hypoxia.

**Fig. 4.**
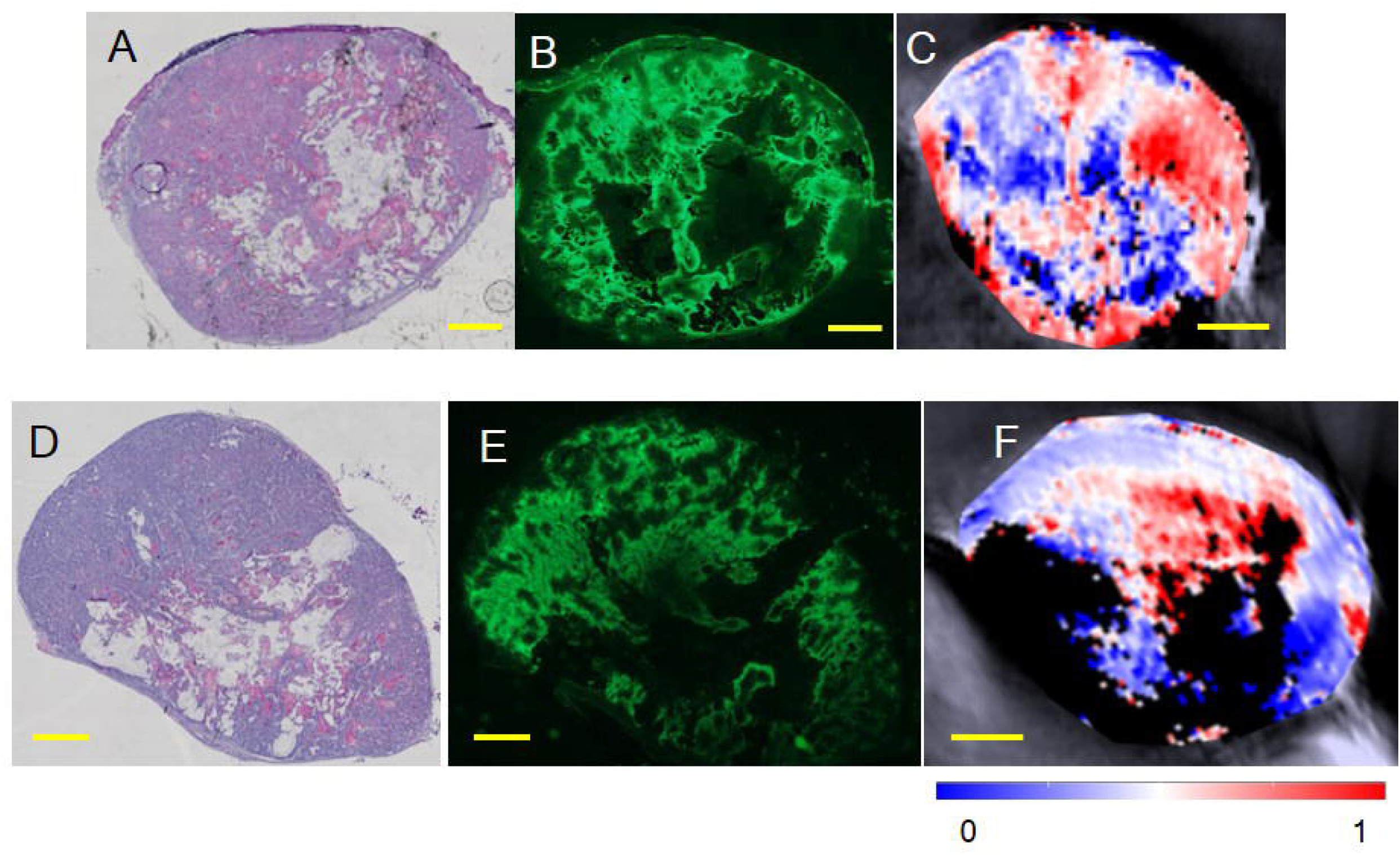
Representative examples of two CAL^R^ tumours (volumes of 220±12 mm^3^, top, and 221±28 mm^3^, bottom). A&D: H&E staining, B&E: pimonidazole staining and C&F: ‘oxymaps’ of corresponding central tumour slices.

### 3.2. Short term PAI variability study

#### 3.2.1. Variation in time over 75 minutes

For the first short-term study, aimed to investigate the anaesthesia effect over 75 minutes in the photoacoustic imaging measurements, four animals were imaged. Fig.5 shows the short term variation in the ROI-averaged percentage of black pixels and average Hb, HbO_2_, HbT and sO_2_, over the three tumour slices per animal. A trend for a decrease in the percentage of black pixels (i.e. those with undetectable levels of haemoglobin), is observed for animals 1, 2 and 4 after the initial 10 minutes of imaging, after an initial increase between 5 and 10 minutes. Animal 3 has a more constant percentage of black pixels compared to the remaining 3 animals, over the 75 minutes of imaging. An increase in Hb and HbO_2_ (and therefore HbT), and sO_2_ over the whole time period was observed for all tumours, except for the sO_2_ values measured for animal 3.

**Fig. 5.**
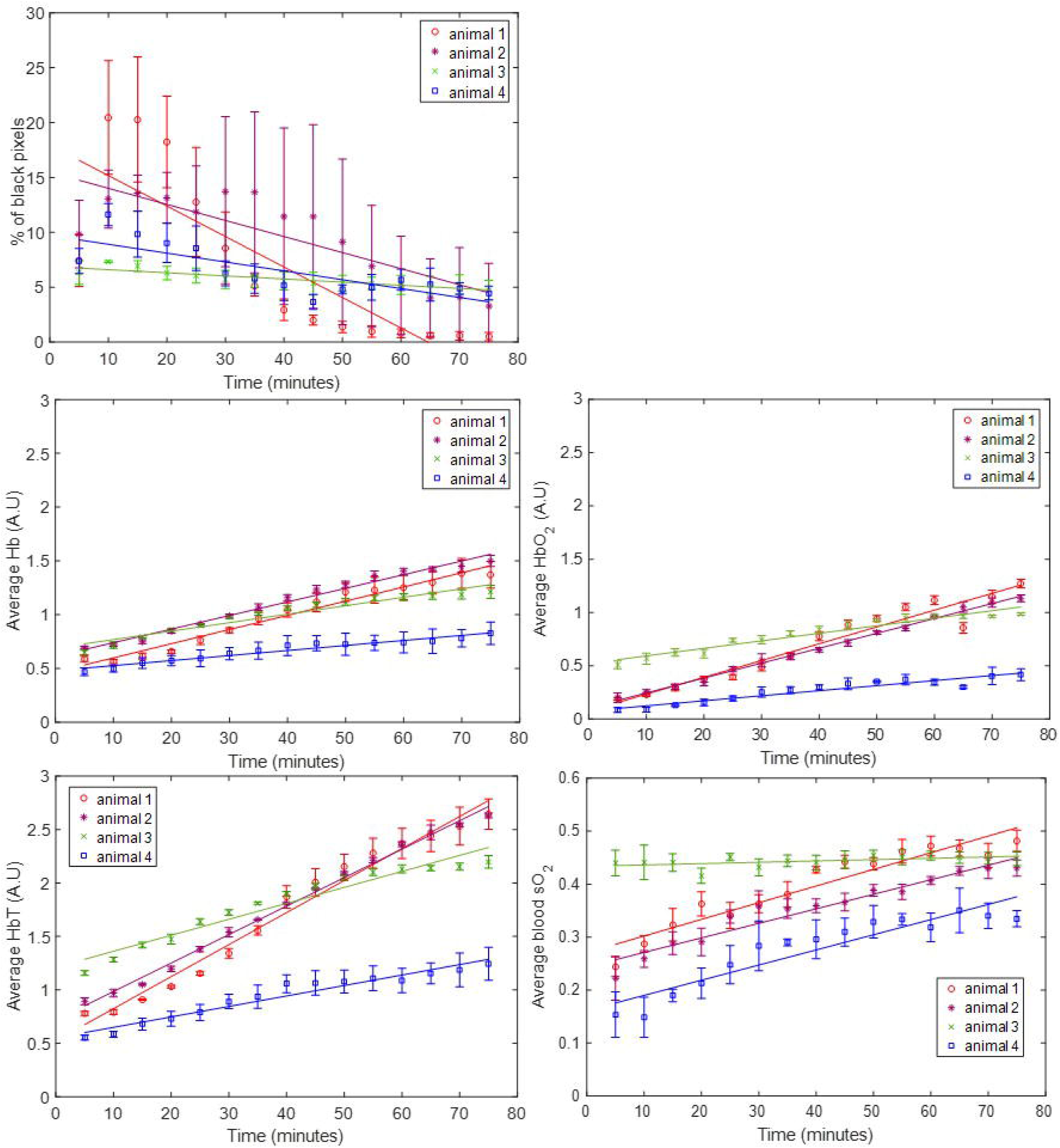
ROI-averaged percentage of black pixels, and Hb, HbO_2_, HbT and sO_2_ signals, for CAL^R^ tumours (n=4) imaged over 75 minutes, during air-breathing. Error bars represent the standard deviation over 3 adjacent central tumour slices (ROIs), 1 mm apart. Solid lines represent a linear regression fitted to the data.

The different change rates are quantified in Table 1. Table 1 summarises the ROI-averaged rate of change of Hb, HbO_2_, HbT, sO_2_ and percentage of black pixels (%BP), as well as the goodness of fit (R^2^) and the sum of squares of error (SSE) of the linear regression used to calculate these rates of signal change. Animals 1 and 2 had a faster increase in Hb (and HbO_2_ than animals 3 and 4. This was reflected in the rate of HbT over 75 minutes. While for animals 1 and 2, the increase in HbT was >0.027 min^−1^, for animals 3 and 4, the increase rate for this parameter was <0.015 min^−1^. Animals 1 and 2 also had a faster decrease in %BP (−0.278 min^−1^ and −0.146 min^−1^, respectively) compared to animals 3 and 4 (−0.0284 min^−1^ and −0.081 min^−1^, respectively).

**Table 1:**
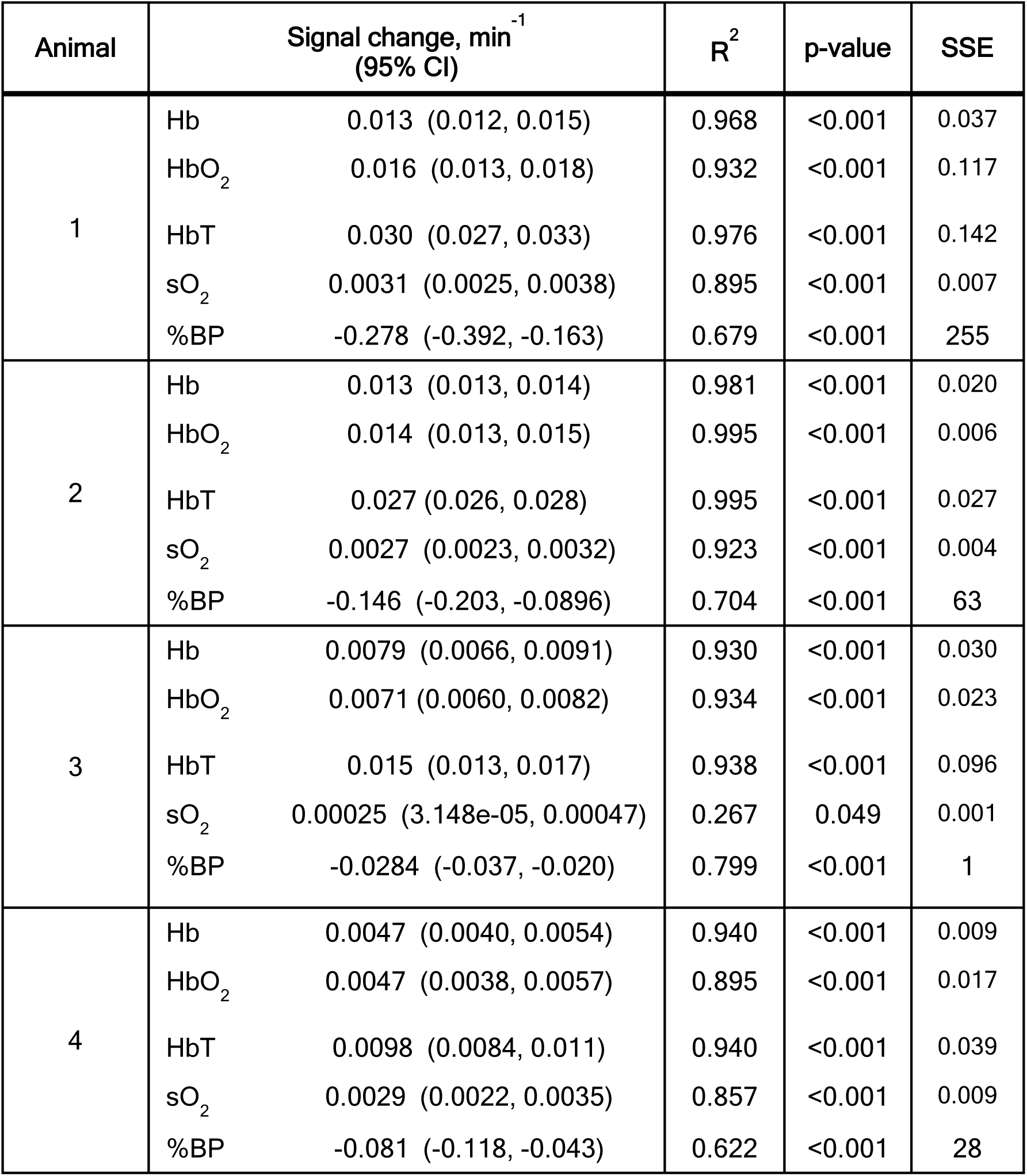
Rate of change of ROI-averaged Hb, HbO_2_, HbT, sO_2_ and percentage of black pixels (%BP) signal during the short-term (75 minutes) longitudinal study. The goodness of fit, R^2^, indicates how much of the signal variability is time dependent and the p-value if the signal change (slope of the linear model) is significantly different from zero. The SSE reflects how similar the predicted values and measured data are, i.e. the closest SSE is to zero, the better the linear fit.

| Animal |  | Signal change, min<sup>-1</sup><br>(95% CI) | R<sup>2</sup> | p-value | SSE |
| --- | --- | --- | --- | --- | --- |
| 1 | Hb | 0.013 (0.012, 0.015) | 0.968 | <0.001 | 0.037 |
|  | HbO<sub>2</sub> | 0.016 (0.013, 0.018) | 0.932 | <0.001 | 0.117 |
|  | HbT | 0.030 (0.027, 0.033) | 0.976 | <0.001 | 0.142 |
|  | sO<sub>2</sub> | 0.0031 (0.0025, 0.0038) | 0.895 | <0.001 | 0.007 |
|  | %BP | -0.278 (-0.392, -0.163) | 0.679 | <0.001 | 255 |
| 2 | Hb | 0.013 (0.013, 0.014) | 0.981 | <0.001 | 0.020 |
|  | HbO<sub>2</sub> | 0.014 (0.013, 0.015) | 0.995 | <0.001 | 0.006 |
|  | HbT | 0.027 (0.026, 0.028) | 0.995 | <0.001 | 0.027 |
|  | sO<sub>2</sub> | 0.0027 (0.0023, 0.0032) | 0.923 | <0.001 | 0.004 |
|  | %BP | -0.146 (-0.203, -0.0896) | 0.704 | <0.001 | 63 |
| 3 | Hb | 0.0079 (0.0066, 0.0091) | 0.930 | <0.001 | 0.030 |
|  | HbO<sub>2</sub> | 0.0071 (0.0060, 0.0082) | 0.934 | <0.001 | 0.023 |
|  | HbT | 0.015 (0.013, 0.017) | 0.938 | <0.001 | 0.096 |
|  | sO<sub>2</sub> | 0.00025 (3.148e-05, 0.00047) | 0.267 | 0.049 | 0.001 |
|  | %BP | -0.0284 (-0.037, -0.020) | 0.799 | <0.001 | 1 |
| 4 | Hb | 0.0047 (0.0040, 0.0054) | 0.940 | <0.001 | 0.009 |
|  | HbO<sub>2</sub> | 0.0047 (0.0038, 0.0057) | 0.895 | <0.001 | 0.017 |
|  | HbT | 0.0098 (0.0084, 0.011) | 0.940 | <0.001 | 0.039 |
|  | sO<sub>2</sub> | 0.0029 (0.0022, 0.0035) | 0.857 | <0.001 | 0.009 |
|  | %BP | -0.081 (-0.118, -0.043) | 0.622 | <0.001 | 28 |

Table 1 also shows the increase in blood sO_2_. Animals 1, 2 and 4, had similar increases in blood sO_2_ (range: 0.0027-0.0031), but for animal 3, the blood sO_2_ rate of change was negligible, 0.00025 (3.148e-05, 0.00047), with R^2^ = 0.267. This is a consequence of a similar increase in Hb (0.0079 min^−1^) and HbO_2_ (0.0071 min^−1^).

The average slice-CoV, intra- and inter-tumour CoVs in ROI-averaged Hb, HbO_2_, HbT and sO_2_, over the 75 minute acquisition for each animal, are summarised in Table S1. Overall, the intra-tumour CoV was higher for the haemoglobin components (Hb: 22.5±6.0%; HbO_2_: 40.2±13.9%; HbT: 22.9±9.2%) than for sO_2_ (16.0±8.4%). The slice-CoV, indicative of the variation of photoacoustic imaging parameters, in the 3 ROIs chosen for each tumour, was lowest for all the parameters (<13.9±7.4%) compared to the intra-tumour CoV.

The inter-tumour CoV calculated for this study was 20.9±3.1% for Hb, 46.4±12.4% for HbO_2_, 28.6±2.3% for HbT and 21.8±10.9% for sO_2_. The largest variation was consistently measured for HbO_2_. These results show that, for the short-term longitudinal study, inter- and intra-tumour CoV were similar, showing a high variability is expected for each animal when imaged over 75 minutes.

#### 3.2.2. Re-positioning study

The results for the study on the effect of repositioning 4 animals three times in the photoacoustic system are shown in Figs. 6 and 7. The trends for air- and oxygen-breathing were similar, so the results for air-breathing are shown here and the results for oxygen-breathing are in the supplementary material section (Fig. S1 and Tables S3-4).

**Fig. 6.**
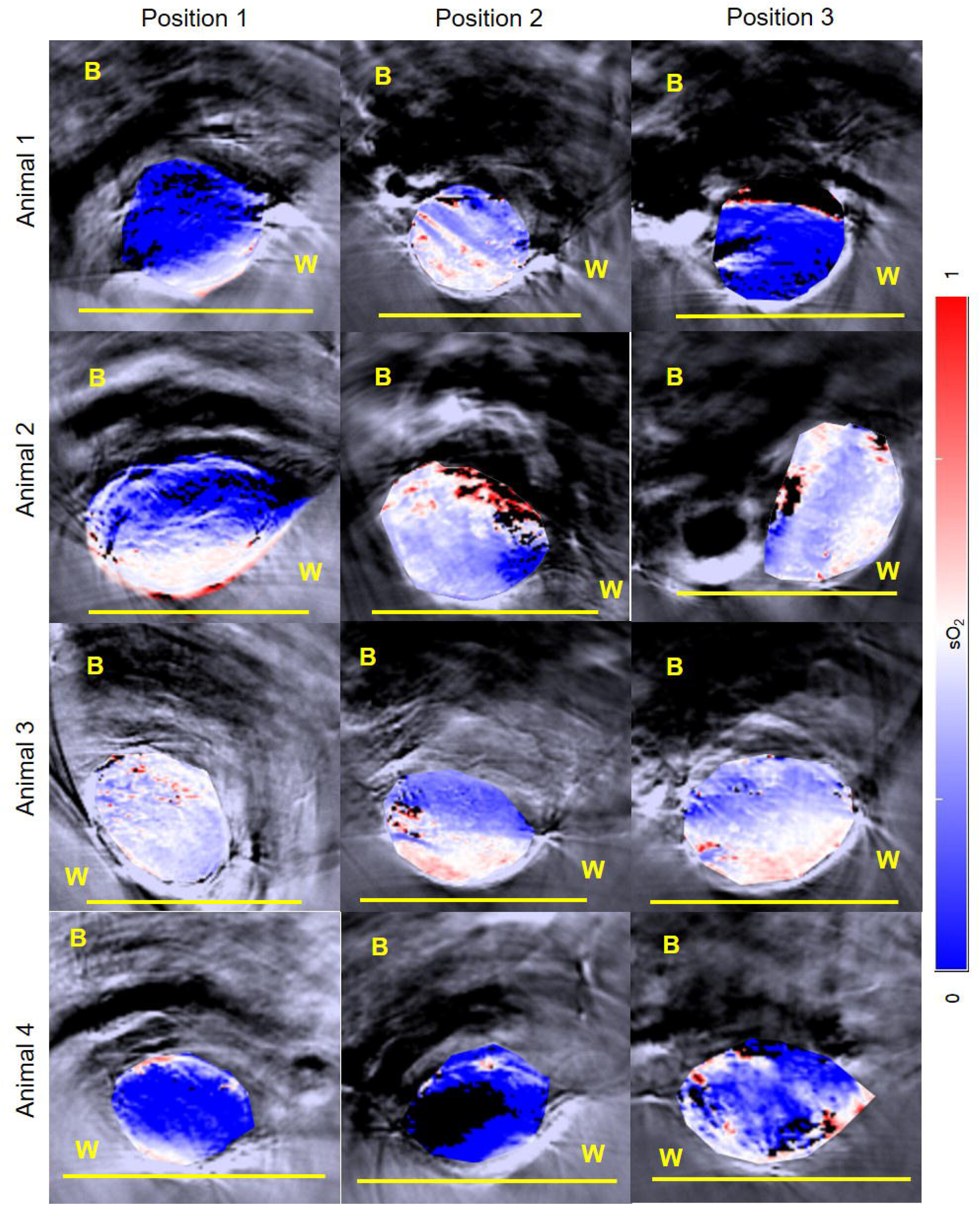
sO_2_ distributions or ‘oxymaps’ of 4 CAL^R^ tumours, positioned differently by remounting the animals 3 times in the MSOT tank with the minimum time interval between each imaging session (see Fig. 2). The ‘oxymaps’ are shown as overlays on greyscale photoacoustic images, of the central slice of each tumour. B: body; W: water Yellow bar represents a 10 mm size scale.

**Fig. 7.**
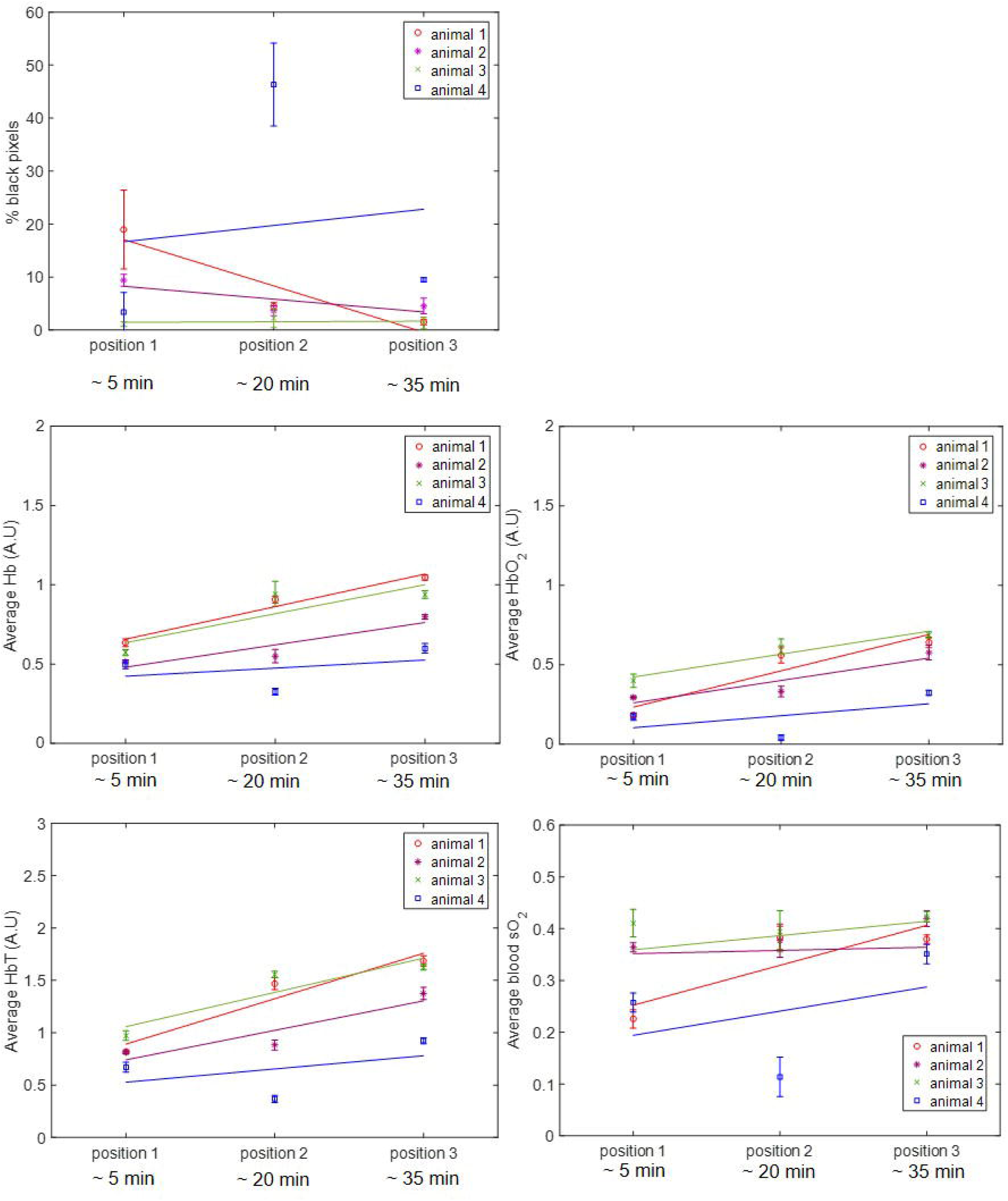
ROI-averaged percentage of black pixel, and Hb, HbO_2_, HbT and sO_2_ signals, for CAL^R^ tumours (n=4) imaged after being removed and re-positioned in the tank three times, during air-breathing. Error bars represent the standard deviation over 3 adjacent central tumour slices, 1 mm apart. Solid lines represent a linear regression fitted to the data.

The ‘oxymap’ of the central tumour slice for each mounting position for each CAL^R^ tumour is shown overlain on greyscale anatomical images in Fig.6. The tumour of animal 1 had mostly low blood sO_2_, <0.5, as indicated by the blue regions (see colour scale), for positions 1 and 3, with the ‘oxymap’ for position 2 indicating some regions with medium to high blood sO_2_ (>0.5), represented in white, pink and red. For animals 2 and 3, the centre of the tumour had mostly low blood sO_2_ (in blue). In the margins of the tumours, white to red pixels are observed, indicative of medium to high blood sO_2_. For animal 3, tumour positions 2 and 3 were similar, as well as the respective ‘oxymaps’. On the bottom row, it is possible to see that the blood sO_2_ of animal 4 is low, and in position 2, the ‘oxymap’ of the central slice of the tumour shows a large region of black pixels (41%). In positions 1 and 3, some white to red pixels are observed, again close to the margins of the tumour.

Following the changes observed in the ‘oxymaps’ in Fig. 6, the temporal changes in the ROI-averaged Hb, HbO_2_, HbT, sO_2_ and percentage of black pixels, were analysed, in order to investigate if re-positioning of animal had an additive effect to the time the animal is under anaesthesia in the variability of the parameters analysed.

Fig.7 shows the ROI-averaged percentage of black pixels for each animal, in each of the three imaging positions, for the 3 central slices. While for the majority of the tumours the mean percentage of black pixels (undetectable haemoglobin levels) was below 19%, animal 4, in position 2, had an exceptionally high proportion of black pixels, 46±8%. Fig. 7 also shows that re-positioning the animals resulted in significantly different measurements of haemoglobin content and blood sO_2_ levels. A trend for an increase in ROI-averaged Hb, HbO_2_ and HbT was observed for all animals over time, independently of the starting and ending angle of the tumour relative to the transducer, apart from animal 4 in position 2, where a high percentage of black pixels was measured. ROI-averaged blood sO_2_ tended to increase for animals 3 and 4 but remain more constant for animals 1 and 2.

The quantification of the changes in signal, both for the individual haemoglobin components, blood sO_2_ and percentage of black pixels, is shown in Table 2. As observed in the 75 minutes imaging study, a trend for an increase in Hb (>0.0034 min^−1^), HbO_2_ (>0.0050 min^−1^) and HbT (>0.0084 min^−1^) was observed over time. Animals 1 and 2 also showed a decrease in %BP (−0.852 min^−1^ and −0.162 min^−1^). For animal 3, since the blood sO_2_ remained constant, R^2^ was low (0.189), although SSE was also very low (<0.001), showing a good linear fit. The R^2^ for the linear regression model applied in the re-positioning study for animal 4 was low for the blood sO_2_ (0.153) and for individual haemoglobin components, <0.289. From Fig. 7, it may be seen that, for this animal, position 2 had abnormally low haemoglobin and sO_2_ values, which may explain the poor goodness of fit. The p-values in Table 2 show that the slopes, i.e. changes in signal over time, for Hb, HbO_2_, HbT and SO_2_, were not significantly different from zero.

**Table 2:** Rate of change of ROI-averaged Hb, HbO_2_, HbT, sO_2_ and percentage of black pixels (%BP) signal during the re-positioning study (air-breathing). The goodness of fit, R^2^, indicates how much of the signal variability is time dependent and the p-value if the signal change (slope of the linear model) is significantly different from zero. The SSE reflects how similar the predicted values and measured data are, i.e. the closest SSE is to zero, the better the linear fit.

| Animal |  | Signal change, min<sup>-1</sup><br>(95% CI) | R<sup>2</sup> | p-value | SSE |
| --- | --- | --- | --- | --- | --- |
| 1 | Hb | 0.014 (-0.019, 0.047) | 0.966 | 0.120 | 0.003 |
|  | HbO<sub>2</sub> | 0.015 (-0.058, 0.089) | 0.874 | 0.230 | 0.015 |
|  | HbT | 0.030 (-0.077, 0.14) | 0.923 | 0.179 | 0.031 |
|  | sO<sub>2</sub> | 0.0051 (-0.034, 0.044) | 0.738 | 0.340 | 0.004 |
|  | %BP | -0.582 (-3.38, 2.21) | 0.875 | 0.231 | 22 |
| 2 | Hb | 0.0094 (-0.043, 0.062) | 0.837 | 0.265 | 0.008 |
|  | HbO<sub>2</sub> | 0.0094 (-0.042, 0.060) | 0.846 | 0.256 | 0.007 |
|  | HbT | 0.019 (-0.085, 0.12) | 0.841 | 0.262 | 0.030 |
|  | sO<sub>2</sub> | 0.0018 (-0.0055, 0.0092) | 0.911 | 0.188 | <0.001 |
|  | %BP | -0.162 (-1.79, 1.47) | 0.612 | 0.428 | 7 |
| 3 | Hb | 0.012 (-0.079, 0.10) | 0.742 | 0.339 | 0.023 |
|  | HbO<sub>2</sub> | 0.0096 (-0.025, 0.044) | 0.927 | 0.174 | 0.003 |
|  | HbT | 0.022 (-0.10, 0.15) | 0.829 | 0.271 | 0.043 |
|  | sO<sub>2</sub> | 0.00041 (-0.010, 0.011) | 0.189 | 0.018 | <0.001 |
|  | %BP | 0.0071 (-0.490, 0.504) | 0.031 | 0.887 | 0.688 |
| 4 | Hb | 0.0034 (-0.11, 0.11) | 0.136 | 0.762 | 0.033 |
|  | HbO<sub>2</sub> | 0.0050 (-0.096, 0.11) | 0.289 | 0.639 | 0.028 |
|  | HbT | 0.0084 (-0.20, 0.22) | 0.207 | 0.700 | 0.122 |
|  | sO<sub>2</sub> | 0.0031 (-0.090, 0.096) | 0.153 | 0.155 | 0.024 |
|  | %BP | 0.204 (-19.3, 19.7) | 0.017 | 0.917 | 1062 |

No significant differences were found between the (temporal) signal change calculated for the repositioning study and that obtained for the longitudinal (75 minutes) study, for ROI-averaged Hb, HbO_2_, HbT and sO_2_, apart for the HbT parameter for animal 2. Figs. S2-S5 show the longitudinal and re-positioning data for Hb, HbO_2_, HbT and SO_2_, for each animal, plotted in the same figure and the p-values showing the similarity of the slopes, i.e. if p-value<0.05 the temporal change in signal was significantly different between the 75 minute and re-positioning studies.

The slice- and intra-tumour CoVs for ROI-averaged Hb, HbO_2_, HbT and sO_2_, are summarised in Table S2. The intra-tumour CoV measurement of animal four, excluding position 2, where 46% of signal was lost, is also shown. As was seen for the 75 minute study, the intra-tumour CoVs for measurements acquired after re-positioning the animals was higher for the haemoglobin components (Hb: 22.0±6.0%; HbO_2_: 40.0±11.0%; HbT: 28.1±5.1%) than for blood sO_2_ (22.0±21.2 and 15.0±11.3 with and without considering position 2 imaging for animal 4, respectively). The slice-CoV was also lower (19.4±18.4% including animal 4, position 2 imaging) compared to the intra-tumour CoV.

Including the data point for animal 4, position 2, the inter-CoV calculated was of 25.8±16.4% for the average Hb, 45.6±19.6% for the average HbO_2_, 30.6±15.6% for the average HbT and 26.5±8.3% for the average sO_2_. Excluding that data point, the inter-CoV calculated was of 20.4±16.4% for the average Hb, 33.2±6.4% for the average HbO_2_, 22.7±6.8% for the average HbT and 13.0±13.2% for the average sO_2_.

### 3.3. Variation during tumour growth, 6 day study

For the long-term study, 10 CAL^R^ tumours were imaged. The ROI-averaged Hb, HbO_2_, HbT and sO_2_ from 3 central slices per tumour, measured during air-breathing, for the long term (6 day) variability study (total of 4 imaging sessions) are shown in Fig. 8. There was no statistically significant persistent increase or decrease in Hb, HbO_2_, HbT, or sO_2_ over the 6 days, with the exception of a statistically significant decrease in Hb, from 1.25±.0.4 A.U. to 0.9±.0.2 A.U., from day 1 to 3. The oxygen-breathing results are similar and are included in the supplementary material section (Fig. S7 and Table S6).

**Fig. 8.**
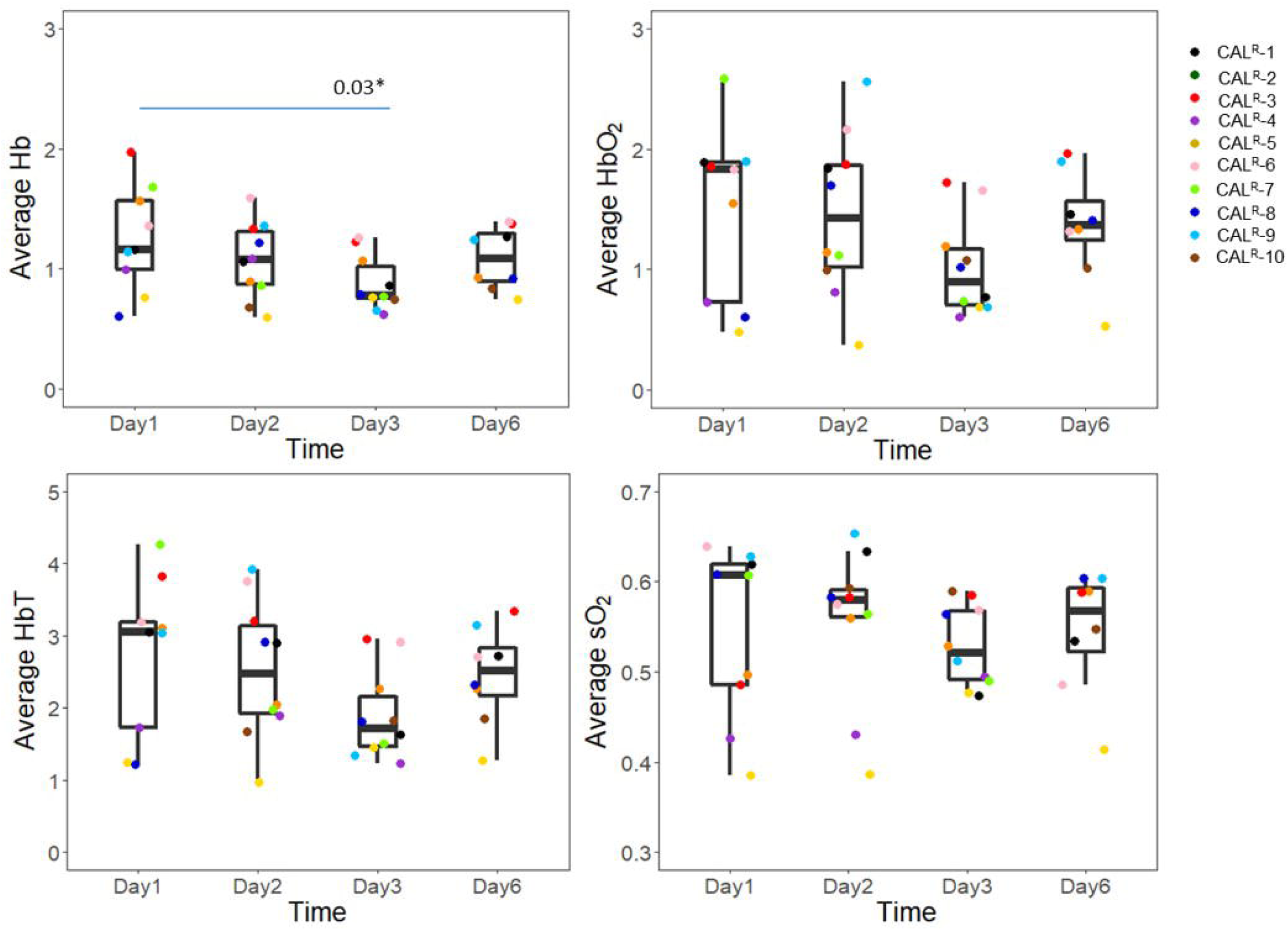
Average haemoglobin and blood sO_2_ values for the 3 central tumour slices of 10 CAL^R^ tumours, over a 6 day imaging period, for air-breathing imaging. Day 1 corresponds to the time at which tumours reached approximately 200 mm ^3^. The whisker plots shows the distribution of the measurements; the central black line in each box represents the median of the population; top and bottom horizontal lines in each box represent the upper and lower quartile and the whiskers the minimum and maximum values.

The inter- and intra-tumour CoV for air-breathing photoacoustic imaging, for 4 imaging sessions acquired over 6 days, are summarised in Table S5. As was found for the short term variability studies, the intra- and inter-tumour CoVs for the sO_2_ parameter were lower (7.5±2.5% and 13.1±3.2%, respectively) than for the haemoglobin parameters (range 19.3±8.7% to 39.7±5.6%).

### 3.4. Tumour growth rate

Growth rates calculated for ten CAL^R^ tumours used in the longitudinal study are shown in Table 3. The individual fitted growth curves, as well as the exponential-linear model fitting, are shown in the supplementary material section (Fig. S6). The uncertainties (95% confidence intervals) are large and hence no statistically significant differences were found between individual tumour growth rates. The goodness of fit, R^2^, for the majority of the tumours (N=8) was ≥ 0.88, suggesting good agreement between the exponential-linear model and the data. Animals CAL^R^-1 and CAL^R^-8, had abnormally low fits to the model. CAL^R^-1 ulcerated, which can affect the tumour growth (see Fig. S6). Fig. S6 shows stagnated tumour growth (108±18 mm^3^ to 123±20 mm^3^) between days 12 to 20 after tumour implantation for CAL^R^-8.

**Table 3:**
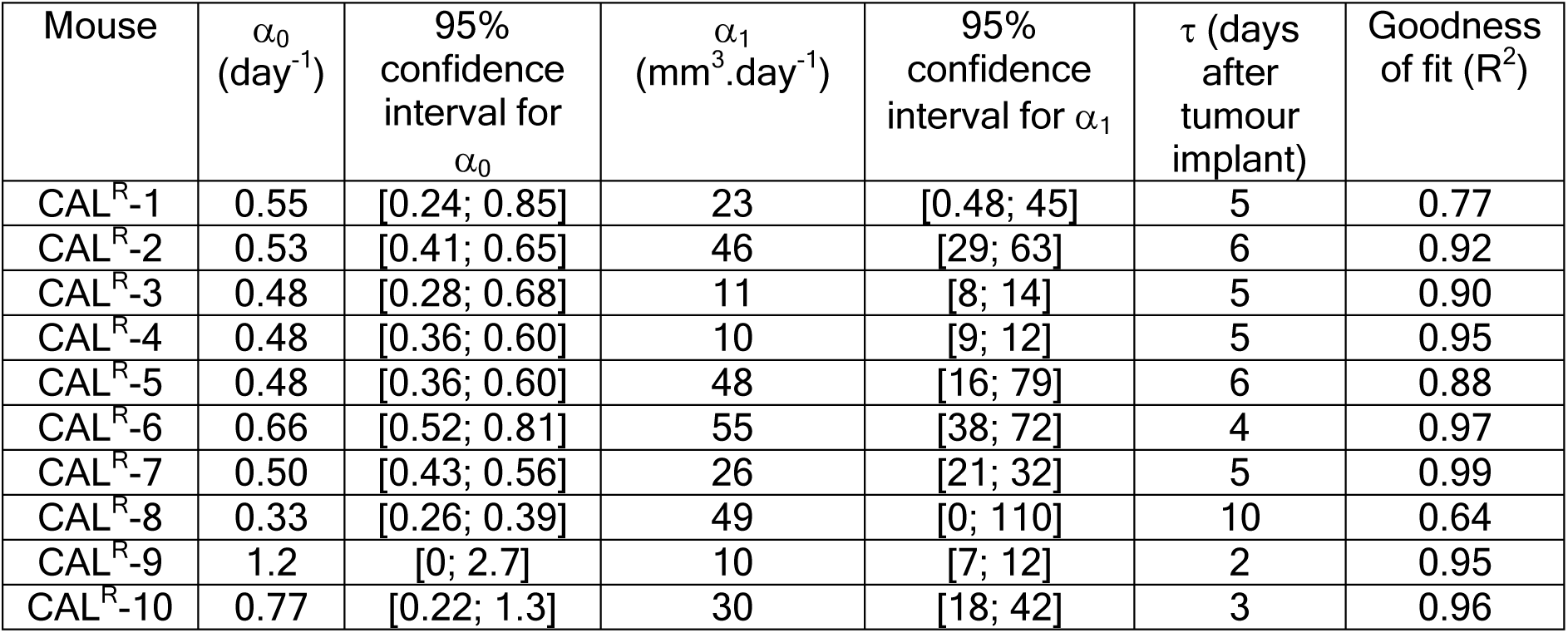
Exponential (α_0_) and linear (α_1_) growth rates, and time after implantation for transition between exponential and linear phase, τ, for 10 CAL^R^ tumours, including the goodness of fit of the exponential-linear model to the growth curves.

Table 6 shows that the time at which tumour growth changed from exponential to linear varied from 2 to 10 days after tumour implant. Based on these results and the growth curves shown in Fig. S6, all photoacoustic imaging acquired for this study was performed during linear growth. In Fig. 8, the change in blood sO_2_ over the 6 days of imaging did not follow the same trend for all tumours, i.e. there was no statistically significant persistent increase or decrease in blood sO_2_ between day 1 and day 6 of imaging. In order to investigate if these changes were related with change in volume over time (ΔV), calculated using raw caliper volume measurements, or linear rate of tumour growth (α_1_), estimated using the exponential-linear model, the relationship between these parameters and the percentage difference in sO_2_ between day 6 and day 1 was investigated and it is shown in Fig. 9. It is possible to observe a good negative correlation between the percentage difference in sO_2_ and both ΔV and α_1,_ but only during oxygen-breathing, i.e. the larger the change in sO_2_ (oxygen-breathing), the faster the tumour was growing.

**Fig. 9.**
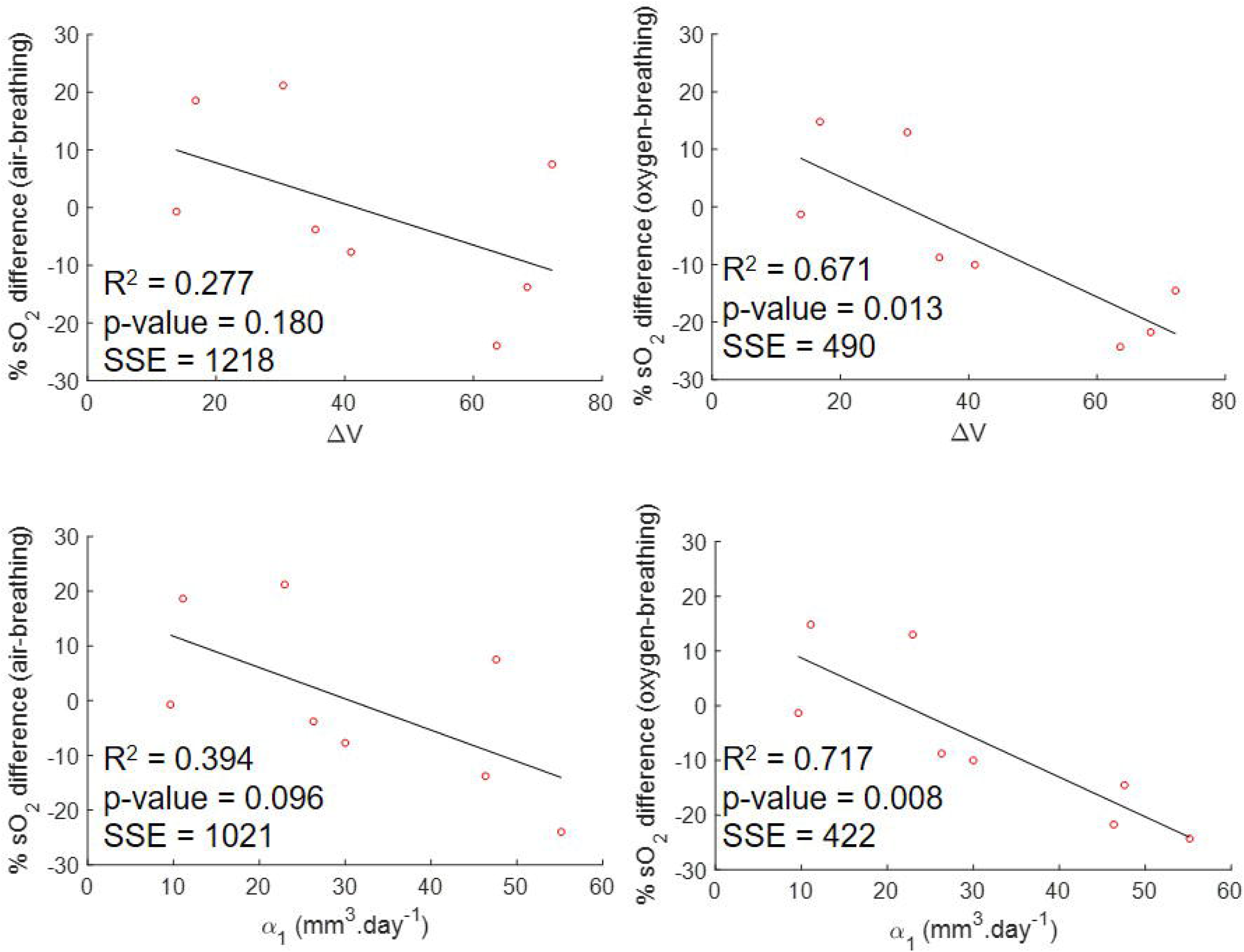
Correlation between rate of change in volume between day 1 and 6 of imaging, ΔV, or linear growth rate, α1, and the percentage difference in sO_2_ between days 1 and 6 of imaging for 8 CAL^R^ tumours, for air- and oxygen-breathing measurements.

## 4. Discussion

The main objective of this paper was to explore the variation of the sO_2_, HbO_2_, Hb and HbT parameters in untreated subcutaneous head and neck tumours measured with a tomographic photoacoustic system, with time over short (minutes-hours) and long periods (several days), and with position of the animals in the imaging cradle. From these studies, it was possible to quantitatively investigate the variability due to anaesthesia, tumour growth and re-positioning. This baseline information will be required for assessing the significance of any changes in these characteristics that may be related to response to treatment, in future studies of the effects of radiotherapy and/or HIFU.

Initially, the spatial correspondence between photoacoustic images and histology staining for hypoxia was investigated (Fig. 4). For this study, pimonidazole, a well-established exogenous marker of hypoxia, was used [37, 38]. Gerling et al. [18] had showed good agreement (R^2^ = 0.915, p-value = 0.0028) between an increased intensity in pimonidazole and low sO_2_ measured using photoacoustic imaging, although with a different tomographic PAI system to that used in this study. In 2016, Tzoumas et al. [39], using the MSOT (iThera) system, also showed good congruence between pimonidazole staining distribution and low blood sO_2_ measurements. The pimonidazole staining from our study, Fig. 4, showed the presence of extensive tumour hypoxia, as has been observed previously using histology and BOLD-MRI with the same tumour model [40]. The pimonidazole distribution was generally in good spatial agreement with blue regions (Fig. 4), i.e., low sO_2_, in the PAI ‘oxymaps’. Fig. 4 also showed that black pixels in the PAI, where the level of haemoglobin falls below the noise threshold of the MSOT, or very dark blue regions (sO_2_^~^0) probably correspond with necrotic regions, as seen in H&E images (A&D). This was expected since regions of depleted of vasculature rapidly become necrotic and acellular. Nevertheless, due to the different slice thicknesses of the techniques (10 µm for histology and 900 µm for PAI), it is not possible to perfectly co-register the two types of images and this may lead to discrepancies. This direct comparison between histology and PAI is also not possible because both modalities are used at different time points. PAI is done *in vivo*, while histology processing is done *ex vivo* and the samples are excised and frozen 45 minutes after imaging, which can contribute to differences between modalities, since hypoxia is variant in time. Despite of the challenges to achieve exact co-registration between histology and MSOT images, the pimonidazole stained tumour sections clearly demonstrated the potential of PAI to infer the spatial heterogeneity in hypoxia in tumours.

The photoacoustic signal variability over 75 minutes was measured to investigate whether the injectable anaesthesia could affect haemoglobin and blood sO_2_ measured in tumours. Fig. 5 shows a decreasing trend of the black pixels, regions where HbT is too low to be measured, after the first 10 minutes of imaging. Shah et al. [23] showed, by co-registration of PAI (using this system) and microbubble contrast-enhanced ultrasound images, that the black regions in tumours (in mice) did not lack perfusion, although typically exhibited lower levels of perfusion and later microbubble arrival than other regions within the tumour. The MSOT and the software available at the moment are still 1^st^ generation and efforts are being made to improve the detection of HbT in order to eliminate the occurrence of black pixels.

During the 75 minutes in which in the percentage of black pixels tended to decrease, an increase in Hb, HbO_2_ and HbT (Fig. 5 and Table 1) was measured, resulting in increased blood sO_2_ for 3 out of 4 animals. These results suggest peripheral vasodilation. The combination of medetomidine, fentanyl and midazolam used for this study has been shown to cause a significant increase in blood pressure in rabbits and rats, which could arise from an increase in vascular resistance or peripheral vasoconstriction, [41, 42]. However, it is well-known that the cardiovascular effects of the combination of these anaesthetic agents vary in different species [43] and to the authors’ knowledge, there have been no reports on its effect in female FOXnu^n1^ mice. The increase in HbT and sO_2_ in our study can be due to a cardiovascular effect of the anaesthetic agent combination; due to an influence of the hydrodynamic pressure, when the animal is enveloped in a plastic film and submerged inside the water tank; or due to an influence of the compression of the tumour due to the animal’s body weight, which possibly stops blood outflow from the tumour. All these factors could have contributed to the trend that was observed and further work is necessary to understand its source.

The trend for an increase in haemoglobin was observed during the whole duration of imaging session, so the signal does not stabilise during an acceptable period of anaesthesia for a small animal. Therefore, this baseline change in signal needs to be considered in future studies, as it is not possible to wait for long enough for a stable signal before starting imaging the animal.

The inter- and intra-tumour CoVs (as shown in Table S1) were similar in this study, >16.0±8.4% and >20.9±3.1%, respectively, regarding the parameters analysed: Hb, HbO_2_, HbT and sO_2_. This reflects a high variation in the measured signal of each animal, expected due to the increasing trend in haemoglobin and blood sO_2_ over time. Joseph et al. (2017) [44] studied the repeatability and reproducibility of PAI with the same system as that used here in the left kidney and spleen of 8–12 week old mice (n=7), acquiring images over a period of 90 minutes, under isoflurane anaesthesia. The authors found a steady increasing trend in HbT and blood sO_2_ over the 90 minute imaging acquisition. The group reported that the intra-organ CoVs for haemoglobin (Hb, HbO_2_ and HbT, measurements averaged over a single ROI) for spleen was below 16.4 and for kidney below 18.5, while for sO_2_ these values dropped to 3.9 and 3.4, respectively. The CoVs measured in this paper during 75 minutes of imaging in subcutaneous tumours were higher than those measured in spleen and kidney, although it is not possible to compare the uncertainties in the measurements. In the experiment conducted by Joseph et al., the organs (spleen and left kidney) were located deep within the animal’s body, so there is probably minimal compression of their blood supply due to mounting in the system, unlike the situation for subcutaneous tumour, located superficially. This is in addition to the deficient vascular supply that renders tumours prone to ‘cyclic’ hypoxia [2], which can affect the measurements of haemoglobin and oxygen saturation.

In the second study, the effects of re-positioning the animal three times during one imaging session were analysed. Fig. 6 shows that the shape of the tumour may vary considerably during repositioning of the animal, changing the size of the tumour-ROI used for analysis. Subcutaneous tumours, besides being subject to compression due to the animal’s body weight acting against buoyancy, as well as due to hydrostatic pressure, can move relative to underlying anatomy as the skin can be highly mobile. Therefore, during longitudinal studies it would be important for the user to place the animal and tumour in the same position relative to the MSOT mouse holder. From Fig.6, it is possible to observe differences in the ‘oxymaps’ depending on the position of the animal, which can result in differences in the distribution and intensity of blue/red pixels. In mouse 4, position 2 resulted in an atypical high number of black pixels (46%), with corresponding uncharacteristically low levels of Hb, HbT and sO_2_. However, this was observed in only one imaging scan, for one animal out of twelve, so the likelihood of this type of artefact is very low.

In order to investigate if the differences in ‘oxymaps’ during the re-positioning study also resulted in significant differences in ROI-averaged photoacoustic imaging measurements (Hb, HbO_2_, HbT and sO_2_) over time, the temporal changes in these measurements were calculated. Fig.7 and Table 2 show a trend for an increase in Hb, HbO_2_, HbT and sO_2_ over time, during the re-positioning study. The signal change reported in Table 2 shows a similar change for animals 1 to 3 for the haemoglobin components (range of 0.0094 to 0.019 for haemoglobin components and of 0.00041 to 0.0051 for sO_2_) compared to that calculated for the 75 minute study (range of 0.0071 to 0.030 for haemoglobin components and of 0.00025 to 0.0031 for sO_2_). A trend for a decrease or constant percentage of black pixels over time, in each position, was also observed for animals 1 and 2, as observed in the 75 minute imaging study. Animal 3 exhibited no trend in levels of sO_2_ compared to the other two animals, consistent with the observations of the sequential imaging for a period of 75 minutes. Table 4 shows that for this study, the p-values were >0.05 so the slopes (change in signal) were not significantly different from zero. This is likely due to the scarce (n=3 positions) number of points, per animal.

From Figs. S2-S5, it is also possible to observe there were no significant differences between the various positions, for the slopes representing the variation over the 75 minutes of the imaging study. The imaging for animal 4, in position 2 that resulted in a large amount of black pixels, increased the inter- and intra-tumour CoV for the re-positioning study. However, the majority of the signal in this imaging scan was lost, so it is not possible to know if this decrease is a result of changes in the tumour or simply an artefact due to the signal loss.

The inter- and intra-tumour CoV for the CAL^R^ tumours due the re-positioning of the animal (Table S2) were >22.0±21.2% and >26.5±8.3%, respectively, for all the parameters (Hb, HbO_2_, HbT and sO_2_), hence similar to those obtained for the study without re-positioning the animal (see Table S1), over a 75 minute period, apart from animal 4. This also suggests that anaesthesia is the main experimental factor that contributes for the temporal variability of the signal, as found by Joseph et al. [44].

The 6 day longitudinal study of CAL^R^ tumours showed no temporal trend in sO_2_ or haemoglobin (Hb, HbO_2_ and HbT) (Fig. 8), i.e. no increase or decrease in these parameters was consistently measured over time for all animals. Tomaszewski et al. [36] used oxygen enhanced (OE)-PAI, to visualise the spatiotemporal heterogeneity of tumour vascular function in a poorly differentiated and aggressive prostate tumour model, PC3. They showed that sO_2_ (while O_2_ breathing) and ΔsO_2_ (difference in measured sO_2_ between air- and oxygen-breathing) decreased with tumour growth, from a 30 mm^3^ volume to time of culling. The CAL^R^ model used for this project has also been shown to develop rapidly [30] and become hypoxic [45], so a decrease in sO_2,_ as observed by Tomaszewski et al. [36], might have been expected. However, this was not the case, either during air-or oxygen-breathing imaging, for the CAL^R^ tumour model.

The intra- and inter-tumour CoVs for the 6 day imaging study were >19.3±8.7% and >27.0±4.1%, respectively, for the haemoglobin parameters (Hb, HbO_2_ and HbT). The sO_2_ parameter showed less intra-tumour CoV (7.5±2.5%) in the long term studies, in comparison to Hb, HbO_2_ and HbT. Joseph et al. [44] also demonstrated, by imaging the animals on 3 consecutive days, that while the sO_2_ was constant over time in kidney (maintained between 0.8 and 0.7) and spleen (maintained between 0.6 and 0.8), for HbT the variation was larger (between 25-16 A.U and 9-13 A.U. for spleen and left kidney, respectively). These studies suggest that variation in blood sO_2_ is expected to be lower compared to that for haemoglobin. Also, the intra-tumour blood sO_2_ variability is less for the long-term study, over 6 days, compared to that obtained for the 75 minute (16.0±8.4%) and re-positioning (22.0±21.2%) studies, once more showing that time under anaesthesia, per imaging session, should be minimised.

In order to assess whether there was any dependence of tumour blood sO_2_ on tumour growth, the growth rates of the 10 CAL^R^ tumours during two different growth stages, were calculated using the exponential-linear model. The goodness of fit showed a good correlation (R^2^>0.8) between the model and 8 datasets. Of the two animals omitted from analysis, one of the tumours became ulcerated, which is known to introduce an error in tumour measurement [46] and the other animal showed an unexpected delay in tumour growth. Fig. 9 shows a good linear relationship between the linear growth rate and the variation in sO_2_ (R^2^ = 0.72, n=8), during oxygen-breathing. A good linear relationship was also found between the rate of change in volume calculated just for the 6 days of imaging and percentage change in sO_2_ (R^2^ = 0.67). Tumours that grew faster (n=5) during the linear phase (α_1_> 20 mm.day^−1^) had a decrease in sO_2_ [range of −10% to −56%] between days 1 and 6 of imaging, probably because they outgrew their blood supply before new vasculature could be formed. Slower growing tumours (n=3) exhibited an increase or no change in sO_2_ [+12% and −1%], possibly because angiogenesis was occurring at a rate similar to the tumour growth rate. In this study, oxygen proved to be a good contrast agent to show the differences between tumours.

A limitation of this study is that it evaluated the baseline variations in the photoacoustic signal for only one tumour type, a subcutaneous head and neck tumour model. Other tumour types might show different variation. For example, Tomaszewski et al. [36] showed that two prostate tumour models, PC3 and LNCaP, exhibited different haemodynamic behaviour, as assessed by PAI.

## 5. Conclusions

This paper documents the variation in haemoglobin (Hb, HbO_2_ and HbT) and oxygen saturation over short (up to 75 minutes) and long (6 days) periods, and the variation with repositioning of the animal, in untreated subcutaneous head and neck tumours in nude mice. Time under anaesthesia is the largest source of temporal variability in the photoacoustic signal. To perform inter- and intra-animal comparison in longitudinal studies, for example when assessing the effects of treatment, ideally the imaging should be started at a fixed time point after anaesthesia and, if possible, sham-treated animals should be imaged at the same time points as the treated animals.

Positioning of the animal was found to influence image quality in a limited number of animals; when reconstruction artefacts or an unexpected absence in PAI signal are observed during the preview of the scan, the animal can be removed and re-positioned to obtain improved PAI signal. With a carefully planned experimental protocol, photoacoustic imaging could be used for assessing changes in tumour haemoglobin and oxygenation in longitudinal studies, including those designed to assess cancer therapies that may affect vasculature and, consequently, tumour oxygenation.

## Acknowledgements

We gratefully thank the Portuguese Foundation for Science and Technology (PhD grant SFRH/BD/90250/2012) for funding part of this project. We further acknowledge funding from the EPSRC (strategic equipment grant EP/NO15266/1) and the Cancer Research UK Cancer Imaging Centre at the Institute of Cancer Research (grant C1060/A16464). We would also like to thank the Focused Ultrasound Foundation for their support of our Centre of Excellence.

